# Single-cell imaging reveals unexpected heterogeneity of TERT expression across cancer cell lines

**DOI:** 10.1101/618769

**Authors:** Teisha J. Rowland, Gabrijela Dumbović, Evan P. Hass, John L. Rinn, Thomas R. Cech

## Abstract

Telomerase is pathologically reactivated in most human cancers, where it maintains chromosomal telomeres and allows immortalization. Because telomerase reverse transcriptase (TERT) is usually the limiting component for telomerase activation, numerous studies have measured TERT mRNA levels in populations of cells or in tissues. However, little is known about TERT expression at the single-cell and single-molecule level. Here we analyzed TERT expression across 10 human cancer lines using single-molecule RNA FISH and made several unexpected findings. First, there was substantial cell-to-cell variation in number of transcription sites and ratio of transcription sites to gene copies. Second, previous classification of lines as having monoallelic or biallelic TERT expression was found to be inadequate for capturing the TERT gene expression patterns. Finally, TERT mRNA had primarily nuclear localization in cancer cells and induced pluripotent stem cells (iPSCs), in stark contrast to the expectation that mature mRNA should be predominantly cytoplasmic. These data reveal unappreciated heterogeneity, complexity, and unconventionality in TERT expression across human cancer cells.

## Introduction

Telomeres, protective structures found at the ends of eukaryotic chromosomes, contain a highly repetitive DNA sequence and associated proteins; they are important for maintaining chromosomal and genomic stability [1]. In early human development, chromosomal telomere shortening that occurs due to the “end-replication problem” during cell proliferation can be compensated by telomerase. Telomerase, first discovered in 1985 in the ciliate *Tetrahymena* [2], is a ribonucleoprotein enzyme that lengthens and maintains the telomeres. After development, telomerase is inactivated in most somatic cells, leading to progressive telomere shortening until a critical length halts cell division and triggers cell senescence (the Hayflick limit). However, telomerase is pathologically reactivated in approximately 80-90% of malignant human cancers, which is considered an early cancer progression event [3–5].

Expression of the catalytic subunit of telomerase, telomerase reverse transcriptase (TERT) [6], is required for telomerase activity. Introduction of TERT expression into normal human somatic cells leads to telomere elongation and cellular immortalization, making its expression necessary, but not sufficient, for driving telomere elongation and oncogenesis in most cancers [7,8]. Increased TERT expression levels have also been found to be associated with poorer patient prognoses for several cancer types, including breast cancer, urothelial bladder carcinomas, non-small cell lung carcinomas, melanoma, and thyroid tumors [9–12], highlighting the importance of understanding the role of TERT expression in cancer and its progression. It is thought that TERT expression, which is relatively low, must be tightly regulated to ensure normal telomere maintenance. Even a small decrease in TERT activity, such as by 10-20%, may potentially result in abnormal telomere maintenance and related pathological conditions (i.e., telomeropathies such as dyskeratosis congenita, aplastic anemia, and pulmonary fibrosis)[13].

While many such TERT expression studies have been published, these studies have been complicated by difficulties in detecting low levels of endogenously expressed TERT mRNA [14] and have typically used methods that look at overall expression levels within a cell line or tissue sample. Traditional cell-population studies do not provide insight into the spatio-temporal aspects of mRNA expression, which has left unanswered – or provided only unclear answers to – many intriguing questions about TERT expression. For example, while population averages can be a good starting point for understanding overall expression levels of a given mRNA, these data cannot answer questions related to expression at the single-cell level, such as how many gene copies are active per cell and whether there is significant heterogeneity between cancer types or among cells in a given cell line, and can even be misleading when such heterogeneity is involved. Additionally, population averages do not identify where molecules of TERT pre-mRNA and spliced (i.e., mature) mRNA reside within cells, even though subcellular localization can have profound effects on the function of RNA molecules.

Recent advancements in RNA imaging [14,15] allow visualization of mRNAs and actively transcribing alleles at the single-cell level even at low abundance. In the current study, we determined single-cell TERT expression levels in several cancer cell lines that had been previously classified as having either monoallelic or biallelic expression (MAE or BAE, respectively) of *TERT* by Huang and colleagues [16]. They determined the MAE or BAE status of these lines by quantifying allelic imbalances of heterozygous single-nucleotide polymorphisms (SNPs) in *TERT* exons using whole-genome sequencing and RNA-sequencing (RNA-Seq) data from the Broad Institute’s Cancer Cell Line Encyclopedia (CCLE) [16]. For reasons that remain unknown, MAE or BAE of *TERT* was found to consistently associate with certain cancer types; for example, melanoma and pancreatic cancer cell lines had MAE of *TERT*, while breast and prostate lines had BAE of *TERT*. Other cancer types were found to be comprised of a mixture of MAE and BAE lines. Overall, 44% (39/88) of cell lines investigated had MAE of *TERT*, while the other lines had BAE of *TERT*. Nearly half (19/39) of MAE lines contained a *TERT* promoter mutation known to reactivate *TERT* expression via transcription factor recruitment [17–20], while all other MAE lines contained no known *TERT* mutations (i.e., apparently “wildtype” lines). It remains unclear how these wildtype lines reactivated *TERT*, or potentially failed to inactivate the gene [21]. Interestingly, Huang and colleagues reported no significant difference in TERT expression levels between mutant and wildtype MAE lines, nor between MAE and BAE lines, though other studies have reported some cancer types that frequently contain *TERT* promoter mutations (e.g., bladder, glioblastoma, and melanoma) to have increased *TERT* expression [12,22,23].

In the current study, we utilized the powerful technique of single-molecule RNA fluorescent *in situ* hybridization (smFISH) to image and analyze individual TERT mRNA molecules and active TERT transcription sites [15]. We found unexpected variance in the number of active transcription sites, both among cells within a given cancer cell line and between different lines, which increased as the mean number of transcription sites in cell lines increased (R^2^=0.92), supportive of transcriptional bursting [24]. *TERT* DNA FISH showed that the number of transcription sites correlated with the number of gene copies (R^2^=0.42), as one might expect. However, we unexpectedly found the MAE and BAE classification of these cancer cell lines to hide much complexity, as the ratio of transcription sites to gene copies generated from our smFISH and DNA FISH data often did not support the cell line’s allelic classification. These data add to our understanding of variance in TERT expression across human cancers, which could help guide future cancer modeling and cancer therapeutic efforts.

## Results

### Validation of TERT single-molecule RNA FISH

To analyze *TERT* active transcription sites at the single-cell level, we employed dual-color smFISH. We designed oligonucleotide probe sets to independently target TERT introns and exons, thus marking the site of transcription (Fig. 1A). To confirm the specificity of the probes, we analyzed their hybridization in HEK293T cells transfected to overexpress (OE) exon-only full-length 3xFLAG-tagged TERT as well as in non-transfected cells. In non-transfected cells, TERT intron signals appeared in the nuclei as punctate dots that co-localized with the TERT exon signals to indicate active transcription sites (Fig. 1B, top panels). HEK293T cells transfected with TERT OE vector showed markedly increased levels of exon probe hybridization (Fig. 1B, middle panels). RNase A treatment eliminated all detectable signal in TERT smFISH (Fig. 1B, bottom panels), confirming that our probes specifically recognize an RNA target. Because not all transfected HEK293T cells overexpressed TERT, we analyzed the co-occurrence of TERT protein (visualized using an anti-FLAG antibody to the 3xFLAG-tagged TERT protein) and TERT mRNA (visualized by TERT exon smFISH). TERT OE cells that were positive for anti-FLAG staining (i.e., expressing TERT protein) also showed clear TERT exon probe hybridization (Fig. 1C, upper panels), while non-transfected cells had much lower levels of mature TERT mRNA and showed no anti-FLAG staining (Fig. 1C, lower panels). These experiments gave considerable confidence that the oligonucleotide probes specifically recognized TERT pre-mRNA and mature mRNA.

**Figure 1.**
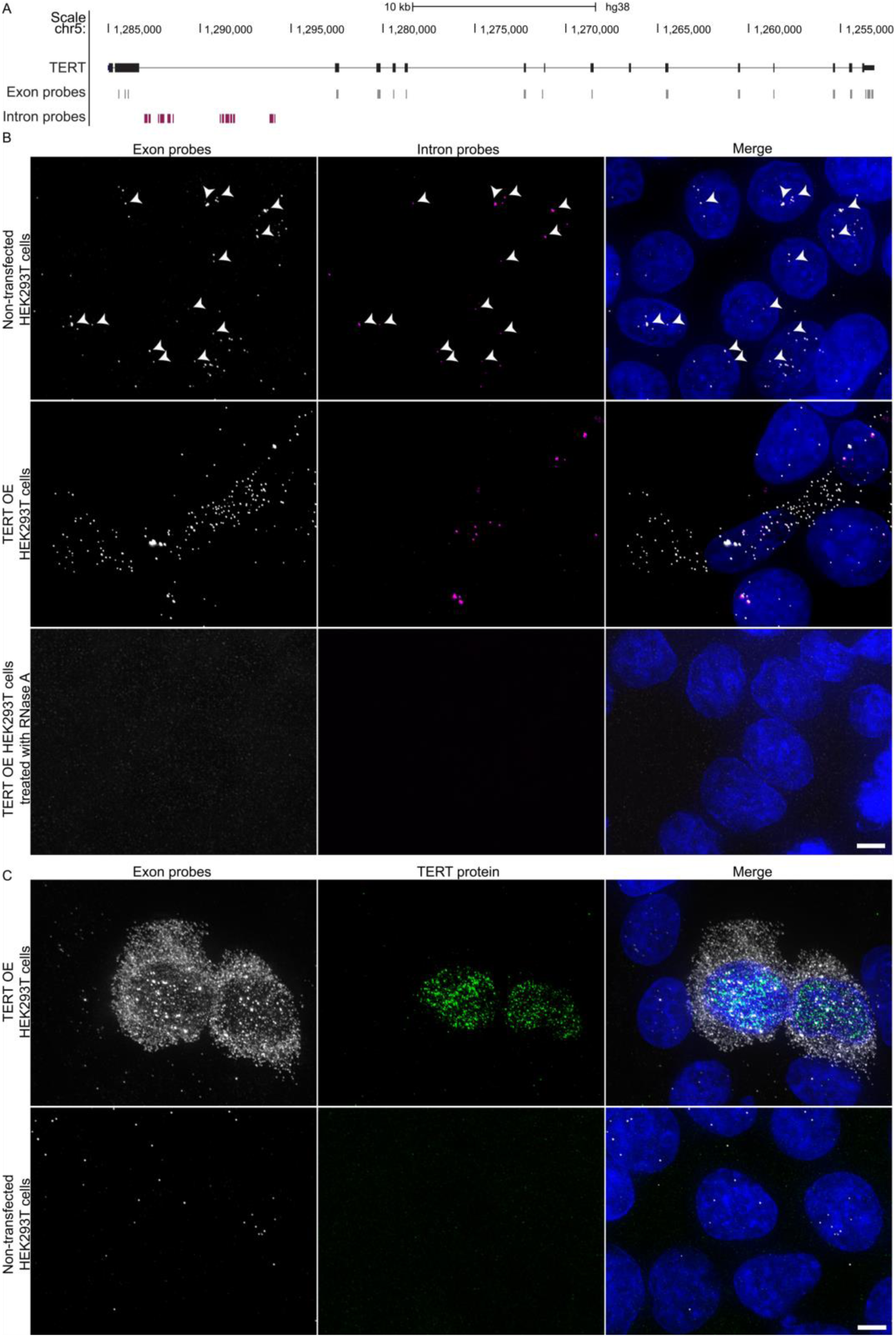
TERT exon and intron single molecule RNA FISH (smFISH) probe design and specificity. (A) The University of California Santa Cruz (UCSC) Genome Browser view showing the localization of *TERT* intron (magenta) and exon (gray) oligonucleotide probes. (B) Maximum intensity projections of TERT exon and intron smFISH of HEK293T cells that were (upper panels) non-transfected, (middle panels) transfected with TERT-3xFLAG over-expression (OE) plasmid, and (lower panels) transfected with TERT OE plasmid treated with RNase A prior to hybridization. Arrowheads indicate representative co-localization of exon and intron signals consistent with active transcription sites. (C) Maximum intensity projections of HEK293T cells that were (upper panels) transfected with TERT-3xFLAG OE plasmid and (lower panels) non-transfected. TERT mRNA was monitored with TERT exon smFISH and TERT protein with immunostaining against the FLAG tag. Data information: TERT exon probes are shown in gray, intron probes in magenta, anti-FLAG immunostaining in green, and DAPI in blue. Scale bars are 5 µm.

As additional controls, we performed GAPDH exon and intron smFISH on TERT OE and non-transfected cells (Fig. EV1). GAPDH exon probes showed characteristic signal in the cytoplasm, and nuclear exon signals co-localizing with intron signals marked active *GAPDH* transcription sites (Fig. EV1B). Both non-transfected and TERT OE transfected cells typically had 0 to 2 exon-intron co-localized nuclear spots. Some HEK293T cells had 3 exon-intron co-localized spots per nucleus, which is not unexpected due to the abnormal karyotype of HEK293T cells [25]. Altogether, these data are supportive of our TERT smFISH probes being specific for detecting and visualizing TERT RNA and *TERT* active transcription sites.

### Unexpected variation of TERT expression across different cancer cell lines

After confirming the specificity of our TERT smFISH probes, we used these probes to visualize active *TERT* genes on a single-cell level across different cancer cell lines. We selected a total of nine cell lines that varied in cancer type and allelic *TERT* expression classification (Table 1). Three of these cancer lines are known to contain a common activating *TERT* promoter mutation (chr5:1,295,228 C>T; hg19), located 124 bp upstream of the *TERT* translation start site (ATG)[16], referred to here as −124 mutants. We also included iPSCs (line WTC-11), which express TERT, and an osteosarcoma cancer cell line (U-2 OS) known to use the alternative lengthening of telomeres (ALT) mechanism and therefore be telomerase-negative[26]. The number of active transcription sites per cell was determined for each cell line based on the co-localization of intron and exon probe signals (Fig. 2A and Table EV1). As expected, in WTC-11 cells the mode number of *TERT* transcription sites was 2 (38% of cells) and the mean was 1.87 (±0.12), while in U-2 OS cells the mode was 0 (97% of cells) and the mean was 0.03 (±0.02). Also, as expected, linear regression analysis across all 11 cell lines tested revealed a strong positive correlation between both the mean number of transcription sites (Fig. 2B) and the TERT RNA levels as measured via qRT-PCR (Fig. 2C) with the mean number of mature mRNA molecules (i.e., the number of exon probe spots per cell) per cell line (R^2^ = 0.64 and 0.63, respectively). GAPDH control smFISH on three cell lines (iPSC line WTC-11 and TERT*-* expressing cancer cell lines SNU-475 and DB) showed expected hybridization patterns with the GAPDH exon and intron probes (Figs. EV3A and EV3B), further supporting the specificity of our TERT smFISH assays. TERT intron and GAPDH intron probes were also used to co-label these cells and gave results similar to those seen when TERT and GAPDH probes were used separately (Fig. EV3C).

**Figure 2.**
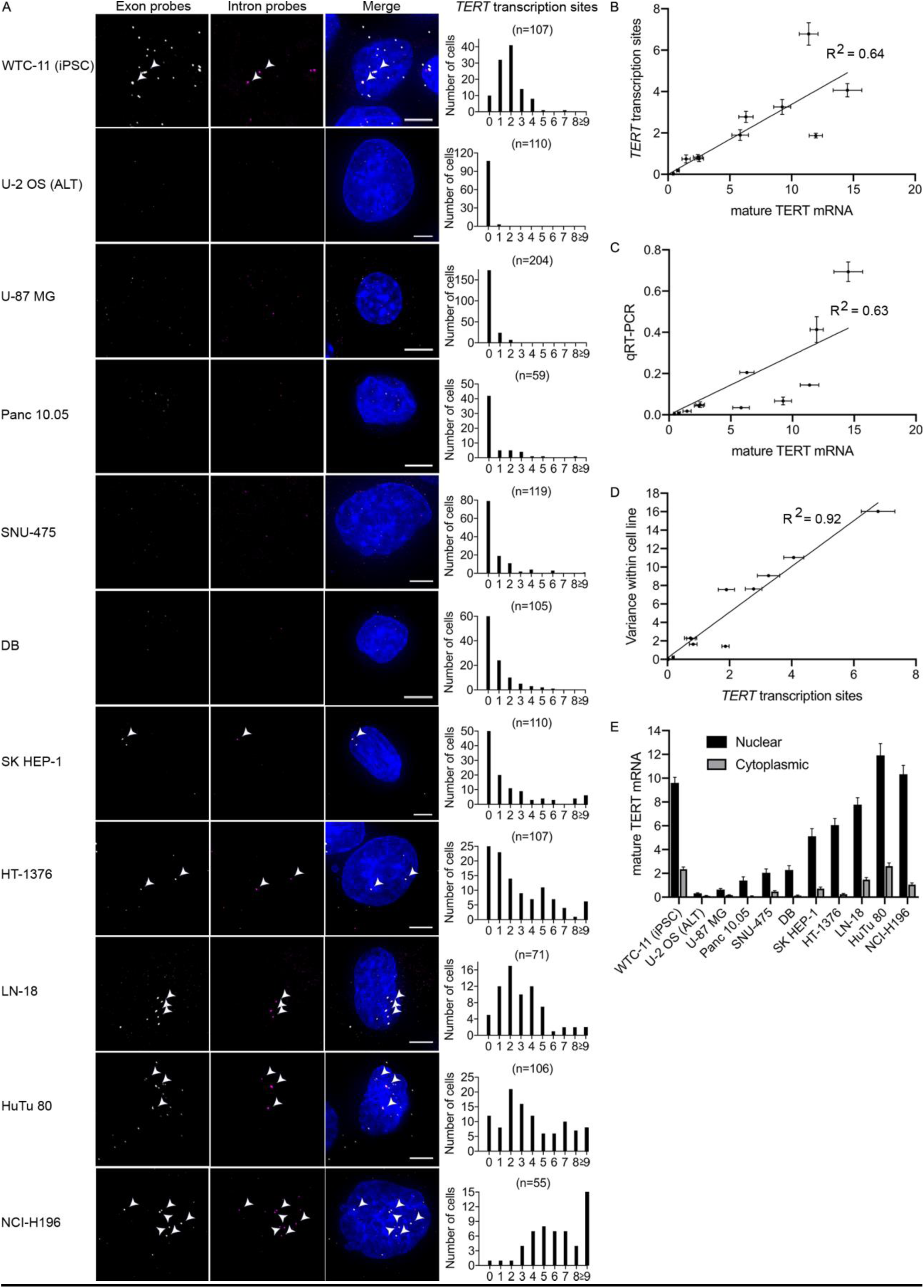
TERT smFISH reveals variation of *TERT* expression across cancer cell lines. (A) Representative single-cell maximum intensity projections of TERT exon and intron smFISH and histograms quantifying cell-to-cell variation in number of transcription sites (number of co-localized exon-intron spots per nucleus) across different cancer cell lines. Arrowheads indicate co-localization of exon and intron signal consistent with active transcription sites. TERT-expressing cancer lines are arranged from least to greatest mean number of active transcription sites (from top to bottom). TERT exon probes are shown in gray, intron probes in magenta, and DAPI in blue. Scale bars are 5 µm. (B) Number of active transcription sites (co-localized exon-intron spots per nucleus measured via smFISH) increases with number of mature mRNA foci (exon spots per cell measured via smFISH). The linear regression line is forced through the 0,0 intersect. (C) Number of mature mRNA foci (exon spots per cell measured via smFISH) correlates with RNA expression levels (measured via qRT-PCR). The linear regression line is forced through the 0,0 intersect. (D) Variance of active transcription sites (co-localized exon-intron spots per nucleus measured via smFISH) within a cell line increases with number of transcription sites. (E) Mature TERT mRNA (exon spots per cell measured via smFISH) primarily has nuclear localization in all lines investigated. Error bars represent standard error of the mean; n = 55 to 204 cells, depending on the cell line, as shown in Fig. 2a. Data information: Error bars represent standard error of the mean. For smFISH, n = 55 to 204 cells, depending on the cell line, as shown in Supplemental Fig. 2a. For qRT-PCR, n = 3 independent measurements.

**Table 1.**
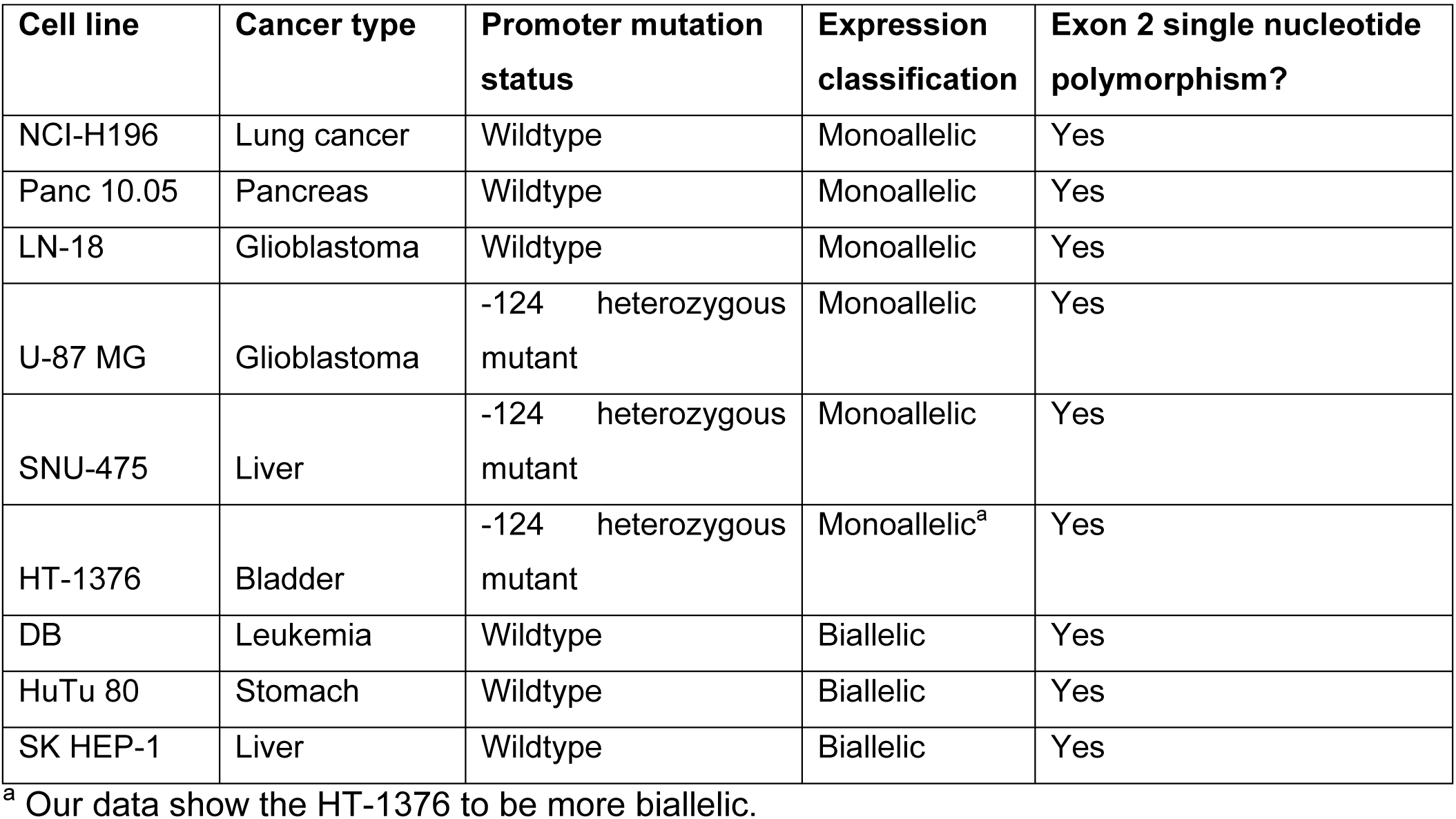
Monoallelic or biallelic *TERT* expression classification and promoter mutation status of TERT-expressing cancer cell lines

The number of active *TERT* transcription sites per cell detected by smFISH varied substantially across the cancer cell lines (see Table EV1 and Fig. 2A for mean and mode values, respectively). The mode number of transcription sites for most telomerase-positive cell lines (6/9 lines) was 0, with only a minority of cells showing one or a few transcription sites. Given that these cancer cells are telomerase-positive and TERT is essential for telomerase activity, this was an unexpected finding (see Discussion). In sharp contrast, other lines had mode values of 2 (LN-18 and HuTu 80) or 5 (NCI-H196), with only a small fraction of the cells showing no transcription sites. Due to variation within the cell lines, while the mean number of transcription sites for most cell lines was less than 1.00 (4/9 lines), mean values for the other cell lines ranged from 1.90 (±0.26) to 6.78 (±0.53). Using linear regression analysis, variance within the 11 cell lines was found to have a strong positive correlation with the number of active transcription sites (R^2^ = 0.92)(Fig. 2D). A similar but weaker correlation was found between variance within the cell lines and the number of mature mRNA molecules (R^2^ = 0.57) (Fig. EV2). It was also surprising to find that while TERT exon probe spots were abundant in the cytoplasm of our HEK293T cells overexpressing TERT (Fig. 1B and 1C), in both iPSCs and cancer cells endogenously expressing TERT, significantly more exon spots were localized within the nucleus instead of the cytoplasm (Fig. 2E; p ≤ 0.0001). Because only cytoplasmic TERT mRNA could be translated into protein, this observation correlates with the low copy number of TERT protein in cancer cells [27]. Overall, our TERT smFISH data indicate that substantial unexpected variation and patterns in TERT expression exists within and across different cancer cell lines.

### Variation in *TERT* gene copy number across different cancer cell lines

To determine whether the variation in TERT expression across different cancer cell lines is due to differences in *TERT* gene copy numbers, *TERT* DNA FISH was performed (Fig 3A). As a positive control, the *TERT* DNA FISH showed the diploid WTC-11 cells to have a mode number of 2 spots per nucleus (94% of cells) and a mean number of 2.01 (±0.04) (see Table EV1 and Fig. EV4 for mean and mode values, respectively). Of the ten cancer cell lines tested, most lines (8/10) had a mode value of 2 to 4 spots per nucleus, with three lines having a mode value of 2 spots (85 – 98% of cells in these lines), four with 3 spots (70 – 90% of cells), and one with 4 spots (86%). In sharp contrast to the smFISH data, these eight cell lines showed a very tight distribution in the number of genes per cell, with low variance within each line. The remaining two cancer cell lines had a mode value of 10 spots per nucleus (24.0 – 35.0% of cells) and relatively higher variance within each line. The mean number of active transcription sites (as determined via TERT smFISH) was found to have a positive correlation with the number of DNA FISH spots across the nine TERT-expressing cancer cell lines (R^2^ = 0.42), a correlation that was dominated by the two high-DNA-copy lines that also had amongst the highest number of transcription sites (Fig. 3B). The mean difference between the number of active sites and DNA FISH spots in the ten cancer cell lines was 2.1 (±0.9) more DNA FISH spots than transcription sites. Overall, these data suggest that the cells are usually not utilizing all copies of the *TERT* gene present.

**Figure 3.**
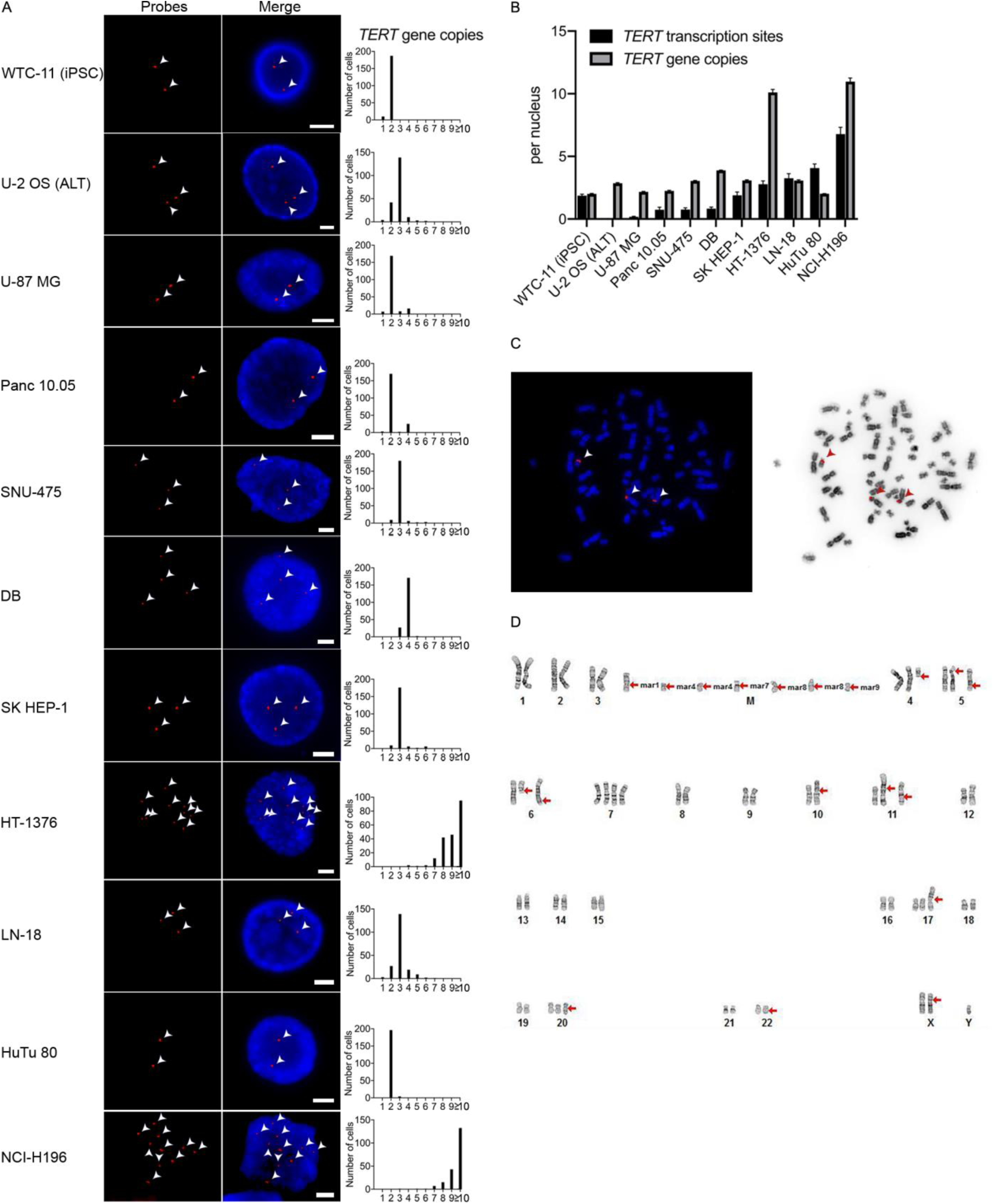
*TERT* DNA FISH reveals more *TERT* genes than active transcription sites in most cancer cell lines. (A) Representative *TERT* DNA FISH single-cell images, with arrowheads indicating probe spots within each nucleus, and histograms quantifying *TERT* DNA FISH, showing variation in *TERT* gene copy number among different cell lines but usually little variation within a given cell line (n = 200 cells for each cell line). TERT-expressing cancer lines are arranged from least to greatest mean number of active transcription sites (from top to bottom). Scale bars are 5 µm. (B) Number of *TERT* gene copies (measured via DNA FISH) increases with the number of active transcription sites (number of co-localized exon-intron spots per nucleus measured via smFISH). Error bars represent standard error of the mean. For smFISH, n = 55 to 204 cells, depending on the cell line, as shown in Fig. 2a. For DNA FISH, n = 200 cells for each cell line. (C) *TERT* DNA FISH of representative metaphase cells from one cancer cell line (LN-18) shows *TERT* triploidy, with arrowheads indicating probe spots, similar to the TERT smFISH and DNA FISH findings for this line. (D) Karyotype analysis of one cancer cell line (LN-18) shows triploidy karyotype, similar to the TERT smFISH and DNA FISH findings for this line. Data information: DNA FISH probes are shown in red and DAPI is shown in blue for A and Cc.

To better understand the genotypic abnormalities observed in the *TERT* DNA FISH, a representative telomerase-expressing cancer cell line (LN-18) was further characterized using *TERT* DNA FISH of metaphase cells and karyotype analysis. The metaphase DNA FISH agreed with the interphase *TERT* DNA FISH performed on this cell line, both showing most cells to have three copies of the *TERT* gene (Fig. 3C). The metaphase analysis additionally showed *TERT* triploidy to be due to amplified full-length and partial copies of chromosome 5, where the *TERT* gene resides. This finding was supported by in-depth karyotype analysis (Fig. 3D), which showed this cell line to have an overall near-triploid (3*n*), unbalanced karyotype with complex abnormalities. Specifically, most cells were XY or XXY and found to contain two to three copies of most chromosomes (with some cells containing four copies of some chromosomes). Regarding chromosome 5, three full or partial copies, sometimes containing unidentifiable material possibly due to chromosomal duplications or deletions, were typically observed in the karyotype analysis. These data indicate that abnormal numbers of the *TERT* gene observed in our cancer lines via *TERT* DNA FISH (i.e., > 2 copies per cell) are not limited to the *TERT* gene, and instead are part of a complex, aneuploid karyotype present in the cancer cells.

### Classification of cancer cell lines as monoallelic or biallelic is insufficient to capture TERT expression patterns

As mentioned earlier, the nine TERT-expressing cancer cell lines used in the present study had been previously classified based on allelic *TERT* expression and *TERT* promoter mutation status (Table 1). To confirm the expected allelic classifications, we used genomic DNA (gDNA) sequencing combined with RT-PCR sequencing of a known SNP in *TERT* exon 2 (rs2736098) (Fig. EV4A). The classification of having either monoallelic or biallelic expression (MAE or BAE, respectively) of *TERT* was confirmed for all nine cell lines, except for the HT-1376 cell line, which was reported to have MAE but appeared to have more BAE (Fig. EV4B; Table 1). While many lines showed similar levels of both alleles based on gDNA sequencing (NCI-H196, Panc 10.05, DB, HuTu 80, and SK HEP-1), several lines had gDNA with relatively greater levels of the inactive allele (LN-18, SNU-475, HT-1376) or active allele (U-87 MG).

Some of the cancer cell lines with apparent MAE of *TERT* had, as expected, roughly half of their *TERT* gene copies active based on our TERT smFISH and DNA FISH data, but this did not necessarily mean there were simply two gene copies with one being active. For example, while Panc 10.05 cells did have on average two (2.25±0.05) *TERT* gene copies and one (0.75±0.20) active transcription site, NCI-H196 cells, surprisingly, had on average 11 (10.98±0.28) gene copies and seven (6.78±0.54) transcription sites (Table EV1). These findings also align with our gDNA sequencing results showing both cell lines to have similar levels of both alleles. Other cell lines with apparent MAE had a greater number of *TERT* gene copies than an expected 2:1 gene copy:transcription site ratio would cause. For example, U-87 MG cells had approximately two (2.17±0.04) gene copies and nearly zero (0.19±0.03) transcription sites and SNU-475 cells had approximately three (3.05±0.04) gene copies and nearly one (0.76±0.14) transcription site. For SNU-475 cells, these findings align with our gDNA and RT-PCR sequencing results showing these cells to have relatively greater levels of gDNA of the inactive allele. Because U-87 MG cells had relatively greater levels of gDNA of the active allele, while only gene copies with the active allele SNP are active, our FISH data suggest that only a fraction of these copies are actually active. Another cell line with apparent MAE, LN-18, had approximately three (3.07±0.06) gene copies and three (3.26±0.36) transcription sites per cell, a 1:1 ratio. These findings are surprising because we also found these cells to have relatively greater levels of gDNA of the inactive allele. Overall, it is noteworthy that while all apparently MAE cell lines only express gene copies with one version of a SNP, there is much underlying complexity, with a ratio of inactive to active gene copies not typically being simply 1:1.

For the cell lines with apparent BAE of *TERT*, both allelic versions of the *TERT* gene would be expected to be active based on our RT-PCR sequencing results. However, our FISH data suggest that three of these four lines have many *TERT* gene copies that are inactive, because there are many more gene copies than active transcription sites. Specifically, SK HEP-1 cells had approximately three (3.09±0.05) gene copies and nearly two (1.90±0.26) transcription sites, DB cells had approximately four (3.88±0.03) gene copies and one (0.83±0.13) transcription site, and HT-1376 cells had approximately 10 (10.11±0.24) gene copies and nearly three (2.78±0.27) transcription sites. For DB cells, there was a slight preference for one allele type to have more active copies than the other, based on our RT-PCR sequencing. This was also seen in HT-1376 cells, though in these cells the active allele was much less common than the inactive allele, based on gDNA sequencing, which aligns with our FISH data showing many more gene copies than transcription sites in these cells. HuTu 80, the other apparently BAE cell line, had approximately two (2.02±0.01) gene copies and four (4.07±0.32) apparent “transcription sites,” which is not possible (see Discussion). For lines HuTu 80 and SK HEP-1, the active copies appeared to be comprised of similar levels of both allele types, based on our RT-PCR sequencing. Overall, for the apparently BAE cell lines, it is interesting that these cells contain many inactive copies of the *TERT* gene (making up most gene copies for half of these cell lines), with the active copies comprising similar levels of both allele versions or a slight preference for one over another.

### Telomere length has little correlation with TERT RNA levels

To determine whether there is a correlation between the number of active *TERT* transcription sites, TERT RNA levels, or *TERT* gene copies and the telomere lengths of these cells, we performed telomere restriction fragment (TRF) analysis (Fig. 4). The iPSCs, ALT cells, and several telomerase-positive cancer cell lines (e.g., DB and HuTu 80) had relatively long telomeres, while other cancer lines (e.g., U-87 MG, Panc 10.05, SK HEP-1, HT-1376, NCI-H196) had relatively short telomeres. Densitometric quantification of the TRF size distributions showed the telomere lengths of the telomerase-positive cancer cell lines and iPSCs (see Table EV1 for cell line mean values) to have little correlation with TERT RNA levels, as measured via qRT-PCR (R^2^ = 0.35; p = 0.07)(Fig. EV5), and no correlation with the number of active TERT transcription sites (measured via *TERT* RNA smFISH; R^2^ = 0.01) or *TERT* gene copies (measured via *TERT* DNA FISH; R^2^ = 0.15). These observations suggest that steady-state telomere length in these cancer lines is ultimately determined by factors beyond TERT expression alone.

**Figure 4.**
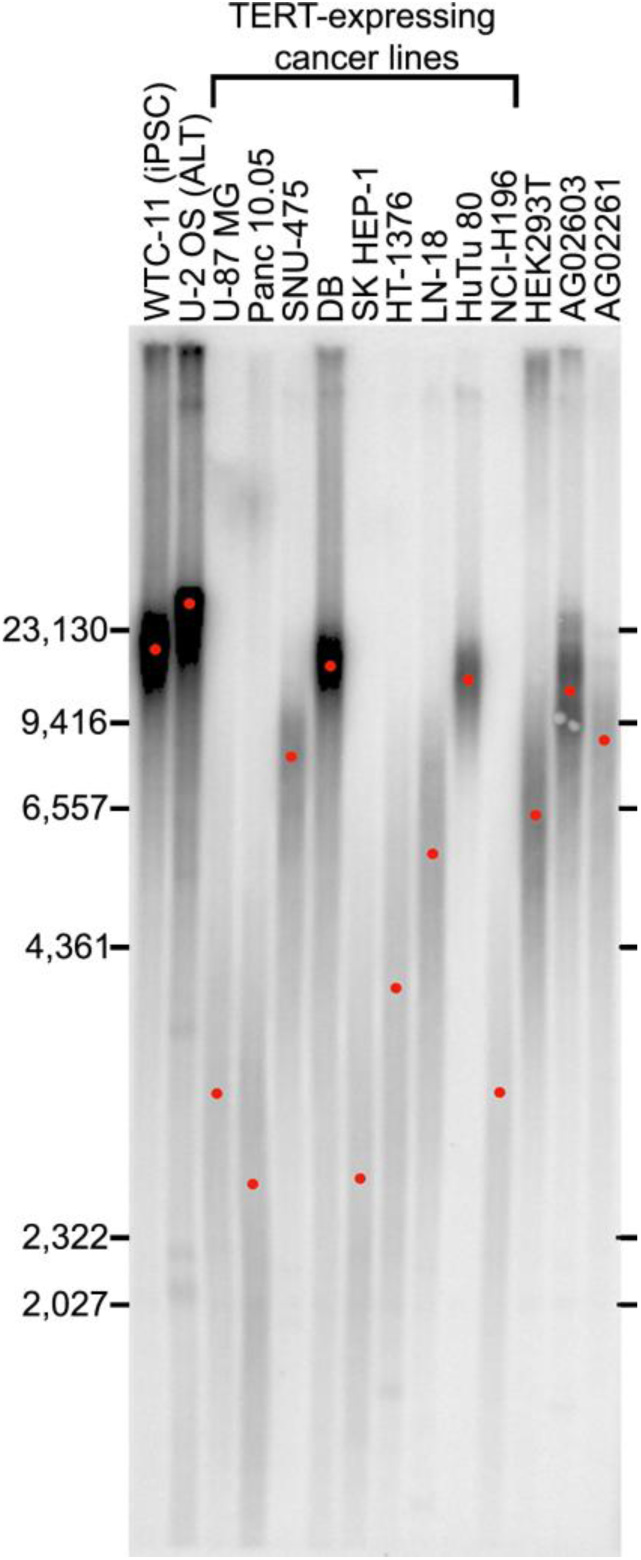
Telomere restriction fragment (TRF) assay of different cell lines, with sizes shown in base pairs (bp) and red dots indicating the mean telomere size for each lane. TERT-expressing cancer lines are arranged from least to greatest mean number of active transcription sites (from left to right).

## Discussion

Utilizing smFISH, we observed great heterogeneity in the number of active *TERT* transcription sites across several different human cancer cell lines and among different cells of a given line. Comparing these data with *TERT* DNA FISH assays, we conclude that previous classifications of cancer cell lines as having monoallelic or biallelic *TERT* expression hide much complexity; the ratios of transcription sites to gene copies in these lines were often unexpected based on their allelic classification.

The variability in single-cell TERT expression levels may reflect the irregular nature of transcription. The correlation we observed between the number of active transcription sites increasing with variance within the cell lines (Fig. 2D) supports transcriptional bursting, or long periods of inactivity interspersed with short periods of strong activity. Specifically, the observed variance is nearly equal to the mean (R^2^=0.92), indicative of a Poisson distribution, which is supportive of transcriptional bursting [24]. We found that nearly all MAE and BAE lines contained many inactive copies of the *TERT* gene, which could also be due to transcriptional bursting and the “snapshot picture” that smFISH allows. We were particularly intrigued to find that six out of nine telomerase-positive cancer cell lines had a modal number of transcription sites of zero. The cell lines were not synchronized, thus some cells might be in phases of the cell cycle when TERT is not actively transcribed. As described above, extremely “bursty” transcription is another possibility; at the extreme, TERT expression might skip an entire cell cycle and then compensate in subsequent cycles. Consistent with our findings, a recent study dedicated to development of TERT RNAscope as a sensitive assay for visualizing TERT mRNA observed a variable pattern of TERT expression in HeLa cells [14]. Taken together, a broad number of telomerase-positive cancer cell lines displays heterogeneity in TERT expression, making it likely that similar heterogeneity in TERT expression is present in primary tumors as well.

Our smFISH experiments also revealed an unexpected subcellular distribution for TERT mRNA. Given that spliced mRNAs are exported from the nucleus for translation, one would expect most mature mRNA to be cytoplasmic. Indeed, this is what we observed for GAPDH mRNA and for overexpressed TERT mRNA (Fig. EV1 and Fig. 1). However, for all nine telomerase-positive cancer cell lines and iPSCs, the mature mRNA (assessed by spots that hybridized only with the exon probes) was mostly nuclear (Fig. 2E). This result is in line with a report by Malhotra et al., 2017 [28], who performed subcellular fractionation of GM12878 and HEK-293 cells and found TERT mRNA to be surprisingly nuclear, as measured by RNA-seq and qRT-PCR. Such a noncanonical distribution of an mRNA could presumably reflect slow export from the nucleus, low cytoplasmic stability, or possibly some nuclear function of TERT mRNA. In any case, the surprising nuclear localization of much TERT mRNA suggests that translatable TERT mRNA copy numbers are even lower than implied by bulk mRNA measurements.

*TERT* DNA FISH showed several lines to have amplified numbers of the *TERT* gene, which was supported by karyotype analysis (Fig. 3D). *TERT* gene copy amplifications are well known, with at least a subset of cells from the following types of cancer cell lines having been reported to have *TERT* gene amplifications: bladder, epidermal, neuroblastomas, hepatic, lung, cervical, breast, colorectal, head and neck, gastrointestinal, osteosarcoma, melanoma, and leukemia carcinomas[10,22,29]. Up to 60 copies have been detected in some individual leukemia cells [29]. Of note, a large study examining 2210 solid tumors spanning 27 tumor tissue types found the 8^th^ most frequent chromosomal gain (13.2% of tumors) to occur in 5p, where the *TERT* gene is located, making the DNA FISH and karyotype findings here not so unexpected [10]. What is new here is the ability to correlate numbers of *TERT* genes with numbers of active transcription sites using smFISH. While we found a general positive correlation (Fig. 3B), we also found that most gene copies are not being actively expressed at any given time.

Because many factors are thought to be involved in determining whether a *TERT* gene is transcribed into RNA, it is not surprising that our observed correlation (R^2^=0.42) was not stronger. For example, genomic rearrangements upstream of, as well as proximal to, *TERT* have been found to cause increased TERT expression in some glioblastomas and neuroblastomas [30,31]. Detecting such subtle rearrangements is beyond the scope of the current study. Cell line-specific variations in the levels of transcription factors that activate or repress *TERT* expression [32] are likely to further complicate correlations between gene copy numbers and transcription levels.

We and others have previously classified cancer cell lines as having MAE or BAE of *TERT* [16,20,33], but our current work reveals that this simple classification is insufficient to capture the complexity of *TERT* gene expression in many cell lines (Table 1, Fig EV4., and Table EV1). While two apparently MAE lines did roughly have the expected 2:1 ratio of *TERT* gene copies to active transcription start sites (based on our FISH data), only one of these (Panc 10.05) had approximately two gene copies and one transcription site, while the other (NCI-H196) had a surprising average 11 gene copies and seven transcription sites (11:7). The three remaining apparently MAE lines (U-87 MG, SNU-475, and LN-18) had unexpected ratios (roughly 2:0, 3:1, and 3:3, respectively). So, while all apparently MAE lines only express gene copies with one version of a SNP, there is much underlying complexity, with a ratio of total gene copies to transcription sites not typically being simply 2:1. Such unexpected ratios could affect studies on TERT that attempt to group these cells primarily based on allelic status and/or promoter mutation status. For the apparently BAE cancer cell lines, both allelic versions of the *TERT* gene are active based on our RT-PCR sequencing results, and thus the active gene copies must be made up of both allele versions. However, our FISH data suggest that these lines rarely have a 2:2 ratio of gene copies to transcription sites, with actual ratios being roughly 3:2, 4:1, and 10:3 for SK HEP-1, DB, and HT-1376 cell lines, respectively.

HuTu80 was the only cell line that showed fewer *TERT* genes than “transcription sites”; its 2:4 ratio should not be possible. Many nuclei contained two very bright spots of intron-exon probe colocalization, which may be the transcription sites, and multiple smaller ones, which may be unspliced TERT pre-mRNA released from the site of transcription. The hypothesis of decreased TERT pre-mRNA splicing efficiency in this particular cell line is testable in the future. Alternatively, we cannot rule out hybridization to RNA from a reactivated pseudogene unique to this cell line, although there have not been reports of a *TERT* pseudogene.

Telomere lengths were found to have an unconvincing correlation with TERT RNA levels via qRT-PCR (Fig. 4 and Fig. EV5; R^2^=0.35; p = 0.07) and did not correlate with TERT active transcription site data via smFISH or *TERT* gene copy numbers via DNA FISH. Previous studies have also not shown a consistent correlation between telomerase activity and *TERT* gene copy number [29]. Such lack of correlation is not surprising, because numerous events separate *TERT* gene copy number from telomerase extension of telomeres. These events include pre-mRNA transcription, splicing, nuclear export, mRNA stability, translation, assembly of the telomerase holoenzyme, and its recruitment to telomeres, each of which may be subject to regulation. For example, our group has previously found that a subpopulation of TERT protein subunits is not assembled into the active telomerase ribonucleoprotein (RNP) enzyme, complicating correlations between TERT expression and telomere length measurements [27]. Finally, telomere length is a balance between extension and shrinkage, so even if TERT RNA levels correlated with extension, they would not necessarily always correlate with steady-state telomere length. Considering these and other complicating factors, it is not surprising that we did not see a stronger correlation between telomere length and metrics of TERT expression or with *TERT* gene copy number.

Overall, the data presented here add to our understanding of heterogeneity in TERT expression across human cancers. We show that TERT expression can be highly variable, both between cancer cell lines and within a given line itself. Being aware of heterogeneity within a cell line may be important for designing effective cancer therapeutics, as subpopulations of cells in a tumor [34] with higher or lower TERT levels may need to be targeted differently to avoid drug tolerance. Our finding that classifying cancer cell lines as simply monoallelic or biallelic does not capture the complexity of their gene expression patterns will also be important when trying to categorize cancers based on differences in TERT expression. Clearly such allelic classifications need to be elaborated upon to properly describe the complex regulation processes at play. Single-cell techniques, such as smFISH and DNA FISH, are clearly powerful tools that may be necessary for proper classification of allelic behaviors, particularly in cancer cells where heterogeneity occurs, as well as for revealing surprising subcellular localizations, such as seen here by the predominantly nuclear localization of mature TERT mRNA.

## Materials and Methods

### Cell lines, culture, and transfection

Lines SNU-475, DB, NCI-H196 (American Type Culture Collection [ATCC]), and Panc 10.05 (University of Colorado Cancer Center, Protein Production/MoAB/Tissue Culture Shared Resource [PPSR]) were maintained in Roswell Park Memorial Institute (RPMI)-1640 medium (Gibco Thermo Fisher Scientific). Lines SK HEP-1, HT-13376, HuTu80 (ATCC), AG02603, and AG02261 (Coriell Institute) were maintained in Eagle’s minimum essential medium (EMEM)(Gibco Thermo Fisher Scientific). Lines U-87 MG (PPSR), LN-18 (ATCC) and U-2 OS (kind gift of David Spector [35]) were maintained in Dulbecco’s modified Eagle medium (DMEM) (Gibco Thermo Fisher Scientific). All media were supplemented with 100 μg/mL penicillin and 100 μg/mL streptomycin (Gibco Thermo Fisher Scientific) and 10% (Sigma-Aldrich), 5% (only line LN-18), or 15% fetal bovine serum (FBS) (AG02603 and AG02261). iPSC line WTC-11 (Coriell Institute) was maintained on recombinant human vitronectin[36,37](Thermo Fisher Scientific), coating six-well tissue culture plastic plates (Thermo Fisher Scientific), with Essential 8 Flex Medium (Thermo Fisher Scientific) and passaged using ethylenediaminetetraacetic acid (EDTA) (Thermo Fisher Scientific). All lines were cultured according to recommended protocols.

For TERT overexpression, 17.5 μg of a plasmid expressing hTERT (kind gift of Joachim Lingner [38]) was transiently transfected into one T-150 flask HEK293T cells using 175 μL lipofectamine 2000 solution (52887, Invitrogen) diluted in OptiMem medium for 4 hours at 37°C. Following the four-hour incubation, media was changed to DMEM supplemented with 6 mM L-glutamine, 10% FBS, 100 μg/mL penicillin, and 100 μg/mL streptomycin. Following overnight culture at 37°C, cells were passaged onto coverglasses prepared for smFISH at a density of 1.7×10^5^ cells/well in 12-well tissue culture plastic plates. Following 48 hours culture at 37°C on the coverglasses, cells were fixed for smFISH.

### Single molecule RNA FISH

Single molecule fluorescent *in situ* hybridization RNA FISH (smFISH) was performed as previously described [15,39]. Tiled oligonucleotides targeting TERT intron and TERT exons labeled with Quasar 570 (TERT intron) and Quasar 670 (TERT exon) were designed with LGC Biosearch Technologies’ Stellaris online RNA FISH probe designer (Stellaris® Probe Designer version 4.2) and produced by LGC Biosearch Technologies. As controls for active transcription detection and proper hybridization to nuclear and cytoplasmic RNAs, we additionally custom designed oligonucleotides targeting GAPDH intron labeled with Quasar 670. GAPDH exon pre-designed probe set [39] labeled with Quasar 670 was purchased from LGC Biosearch Technologies.

Cells were seeded on glass coverslips coated with poly-L-lysine (10 μg/mL in PBS). Before hybridization, coverslips were washed 2x with PBS, fixed in 3.7% formaldehyde in PBS for 10 min at room temperature (RT), followed by 2x wash with PBS. Coverslips were immersed in 70% EtOH and incubated at 4°C for a minimum of 1 h. Coverslips were then washed with 2 mL of Wash buffer A (LGC Biosearch Technologies) at RT for 5 min. RNase A treated controls were incubated in 2 mL of RNase A (200 μg/mL in PBS) for 1 h at 37°C prior washing with wash buffer A. Cells were hybridized with 80 μL hybridization buffer (LGC Biosearch Technologies) containing properly diluted smRNA FISH probes (1:100) overnight at 37°C in a humid chamber. Next day, cells were washed with 1 mL of wash buffer A for 30 min at 37°C, followed by another wash with wash buffer A containing Hoechst DNA stain (1:1000, Thermo Fisher Scientific) for 30 min at 37°C. Coverslips were washed with 1 mL of wash buffer B (LGC Biosearch Technologies) for 5 min at RT, mounted with ProlongGold (Life Technologies) on a glass slide and left to curate overnight at 4°C before proceeding to image acquisition (see below). All smFISH graphs were prepared using GraphPad Prism 8 software (version 8.1.0).

### Single molecule RNA FISH/anti-FLAG immunofluorescence

Coverslips intended for anti-FLAG immunofluorescence and smFISH were processed in the same way as described above, with the following changes: (i) hybridization buffer contained 1:100 dilution of TERT exon and intron probes and 1:800 dilution of primary antibody (mouse M2 monoclonal anti-FLAG, F1804, Sigma), and (ii) the first wash with wash buffer A after ON hybridization contained 1:800 diluted anti-mouse secondary antibody labeled with Alexa Fluor 488 (Abcam, ab150113).

### Microscopy and image analysis

Z-stacks (200 nm z-step) capturing entire cell volume were acquired with a GE wide-field DeltaVision Elite microscope with an Olympus UPlanSApo 100x/1.40-NA Oil Objective lens and a PCO Edge sCMOS camera using corresponding filters. 3D stacks were deconvolved using the built-in DeltaVision SoftWoRx Imaging software. Maximum intensity projections of each image were subjected for quantification using Fiji.

### DNA isolation, polymerase chain reaction (PCR), and sequencing

Genomic DNA (gDNA) was isolated from cells using Quick-DNA Miniprep Kit (11-317AC, Zymo Research). 20 μL PCR reactions were performed using 50 ng gDNA and Phusion High-Fidelity DNA Polymerase (F-530, Thermo Fisher Scientific) supplemented with 7-deaza-2’-deoxy-guanosine-5’-triphosphate (7-Deaza-dGTP)(10988537001, Sigma-Aldrich) to aid in amplifying GC rich regions. Sequences for primers (Integrated DNA Technologies [IDT]) are listed in Table EV2. PCR products were purified using E.Z.N.A. Cycle Pure Kit (D6492, Omega Bio-Tek) and underwent Sanger sequencing (GENEWIZ).

### RNA extraction, cDNA synthesis, and quantitative real-time PCR (qRT-PCR)

Total RNA was isolated from cells using the E.Z.N.A. Total RNA Kit I (R6834, Omega Bio-Tek) and the RNase-free DNase Set I (E1091-02, Omega Bio-Tek) to eliminate potentially contaminating DNA. 1 μg RNA was used to synthesize cDNA, using the SuperScript IV First-Strand Synthesis System (Invitrogen Thermo Fisher Scientific, 18091050) with random hexamers and RNase H treatment. qRT-PCR was performed with SYBR Select Master Mix (4472908, Thermo Fisher Scientific) supplemented with 7-Deaza-dGTP using the LightCycler 480 software (Roche).

Primers used were previously described [12,40], except for *TERT* exon 2 primers (primer sequences are listed in Table EV2). Primer specificity was confirmed using gel electrophoresis, melting temperature analysis, and Sanger sequencing. 10 μL qRT-PCR reactions were run in triplicate on a 96-well plate, and data normalized to the geometric mean of three “housekeeping” genes (glyceraldehyde phosphate dehydrogenase [*GPI*], glucose phosphate isomerase [*PPIA*], and hydroxymethylbilane synthase [*HMBS*]). PCR products to be sequenced were purified using E.Z.N.A. Cycle Pure Kit (D6492, Omega Bio-Tek) and underwent Sanger sequencing (GENEWIZ).

### DNA FISH and karyotype analysis

DNA FISH (Empire Genomics, #TERT-20-OR) of all cell lines and karyotyping of LN-18 cells was performed by the WiCell Research Institute Characterization Laboratory.

### Telomere length by Southern blotting

Telomere restriction fragment length analysis was carried out as previously described [41–44]. Briefly, genomic DNA (gDNA) was isolated from cells using Quick-DNA Miniprep Kit (11-317AC, Zymo Research). 1.5-4.5 µg of gDNA from each cell line was digested with RsaI and HinfI. Digested gDNA samples were resolved on a 0.8% agarose gel. The DNA was then transferred to Hybond N+ Nylon membrane (GE), which was probed for telomeric sequence using a radiolabeled (TTAGGG)_4_ probe. The membrane was imaged using phosphor screens and a Typhoon FLA 9500 Variable Mode Imager (GE)[45].

To calculate mean telomere length, lane intensity profiles were extracted and their centers were found using ImageQuant TL, and lengths of these center points were calculated using a λ-HindIII molecular weight marker (NEB). Graphs were prepared using GraphPad Prism 8 software (version 8.1.0).

## Acknowledgements

We thank Andrew J. Bonham (Metropolitan State University of Denver) and Taeyoung Hwang (University of Colorado Boulder) for thoughtful discussions on this work. We thank Cech lab members Dan Youmans, Yicheng Long, Josh Stern, Ci Ji Lim, and Anne Gooding for useful discussions. We thank Arthur Zaug (Cech lab) for assistance with the TERT overexpression and TRF experiments. We thank Roy Parker and Carolyn Decker (University of Colorado Boulder) for access to, and training on, the DeltaVision Elite microscope. We thank the BioFrontiers Advanced Light Microscopy Core and Joe Dragavon (University of Colorado Boulder) for access to image analysis software. We thank Theresa Nahreini and Nicole Kethley for use of the Cell Culture Facility (University of Colorado Boulder). We thank the WiCell Research Institute Characterization Laboratory, specifically Kim Leonhard and Erik McIntire, for DNA FISH and karyotype analysis and thoughtful discussions. This work was funded by National Institutes of Health grant R01 GM099705 to T.R.C. T.R.C. is an investigator and J.L.R. is a faculty scholar of the Howard Hughes Medical Institute.

## Author contributions

T.J.R. performed cell culture, qRT-PCR and gDNA sequencing experiments, and data analysis; G.D. conducted the smFISH assays; G.D. and T.J.R. designed, imaged and interpreted the smFISH assays; E.P.H. designed and performed the TRF assay; J.L.R. and T.R.C. contributed to overall experimental conception and design and supervised experiments; T.J.R. wrote the manuscript with substantial contributions from all other authors.

## Expanded View figures

**Table EV1.** FISH and qRT-PCR mean values for all cell lines tested

| Cell line | TERT smFISH transcription sites (# co-localized exon-intron spots per nucleus) | TERT smFISH transcription sites variance | TERT smFISH mature mRNA (# exon spots per cell) | TERT qRT-PCR normalized values | GAPDH smFISH transcription sites | GAPDH smFISH transcription sites variance | *TERT* DNA FISH (# spots per nucleus) | Telomere length (bp) |
| --- | --- | --- | --- | --- | --- | --- | --- | --- |
| WTC-11 (iPSC) | 1.87 | 1.42 | 11.96 | 0.41321 | 1.61 | 0.59 | 2.01 | 18354 |
| U-2 OS (ALT) | 0.03 | 0.03 | 0.43 | 0.00552 |  |  | 2.86 | 26814 |
| U-87 MG | 0.19 | 0.22 | 0.80 | 0.00847 |  |  | 2.17 | 2844 |
| Panc 10.05 | 0.75 | 2.30 | 1.46 | 0.01785 |  |  | 2.25 | 2742 |
| SNU-475 | 0.76 | 2.25 | 2.52 | 0.04869 | 1.67 | 3.08 | 3.05 | 8045 |
| DB | 0.83 | 1.64 | 2.42 | 0.04406 | 1.17 | 2.26 | 3.88 | 16551 |
| SK HEP-1 | 1.90 | 7.56 | 5.84 | 0.03439 |  |  | 3.09 | 2745 |
| HT-1376 | 2.78 | 7.63 | 6.31 | 0.20529 |  |  | 10.11 | 3125 |
| LN-18 | 3.26 | 9.06 | 9.24 | 0.06695 |  |  | 3.07 | 5357 |
| HuTu 80 | 4.07 | 11.03 | 14.52 | 0.69334 |  |  | 2.02 | 14809 |
| NCI-H196 | 6.78 | 16.03 | 11.38 | 0.14444 |  |  | 10.98 | 2822 |

**Table EV2.**
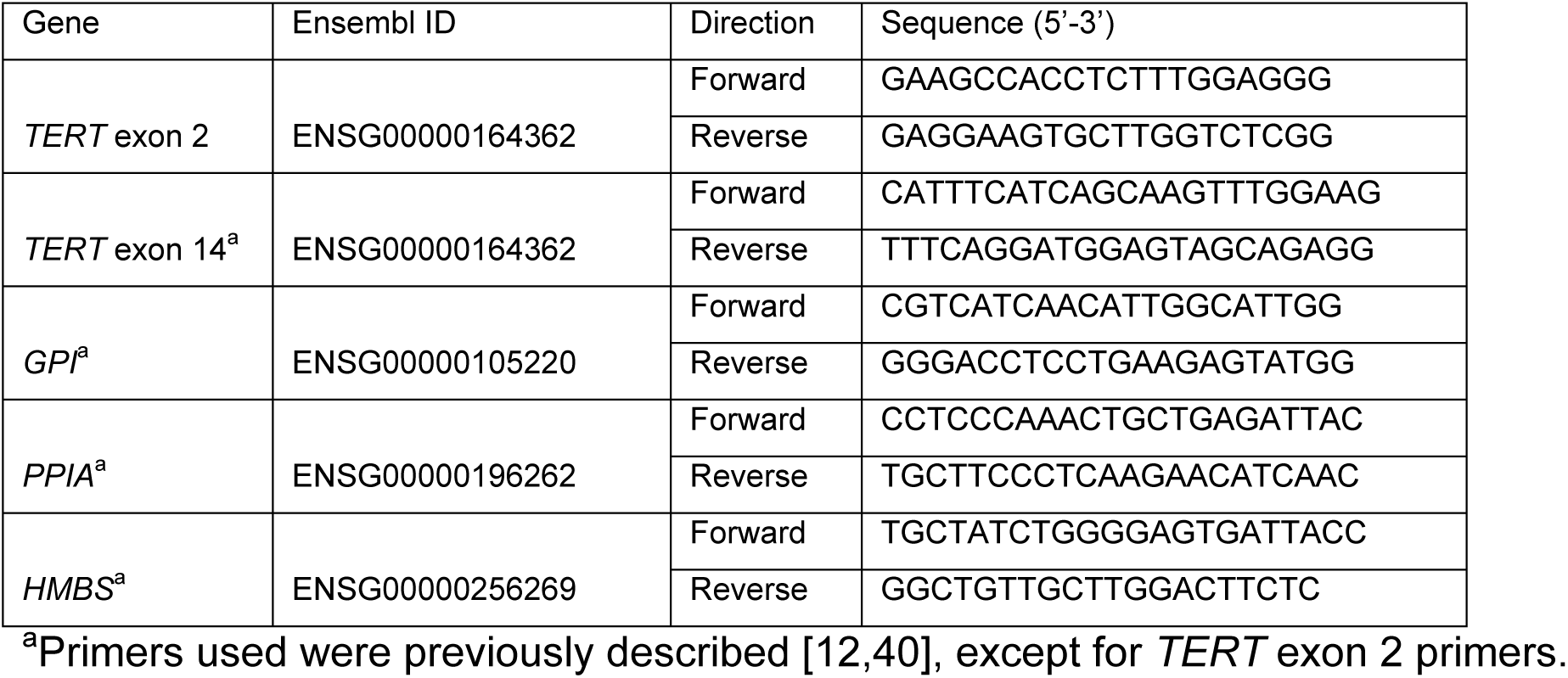
Primer sequences

**Figure EV1.**
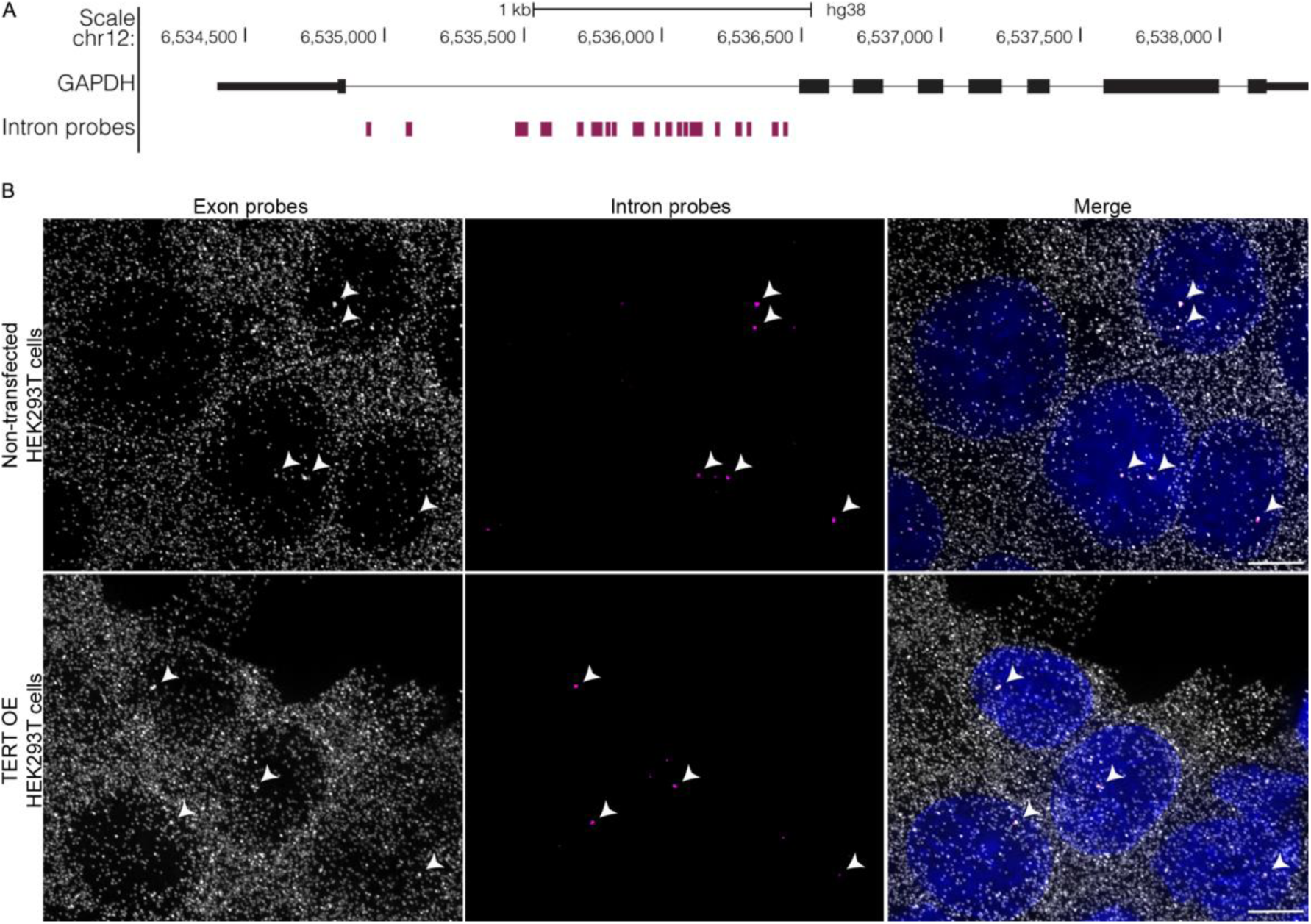
GAPDH exon and intron single molecule RNA FISH (smFISH) design and specificity. (A) The University of California Santa Cruz (UCSC) Genome Browser view showing the localization of GAPDH intron (magenta) oligonucleotide probes. The exon probes were published previously (see Methods). (B) Maximum intensity projections of GAPDH exon and intron smFISH of HEK293T cells that are (upper panels) non-transfected and (lower panels) transfected with TERT-3xFLAG over-expression (OE) plasmid. Data information: Arrowheads indicate co-localization of exon and intron signal consistent with active transcription sites. GAPDH exon probes are shown in gray, intron probes in magenta, and DAPI in blue. Scale bar is 5 µm.

**Figure EV2.**
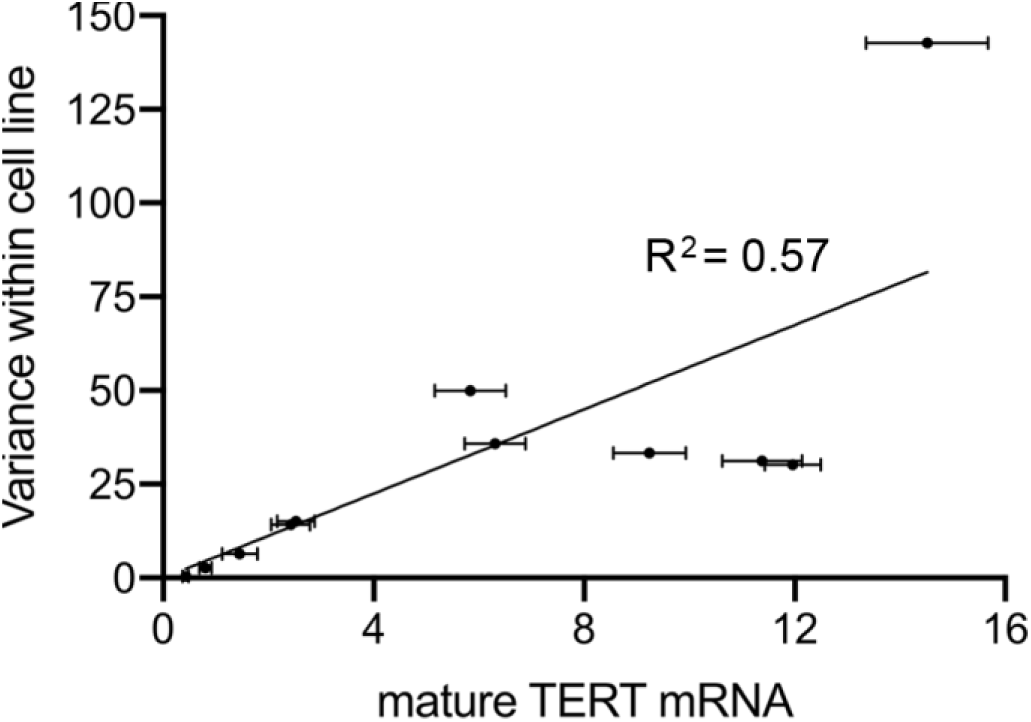
Variance in levels of mature TERT mRNA (exon spots per cell as measured via smFISH) within a cell line increases with these levels.

**Figure EV3.**
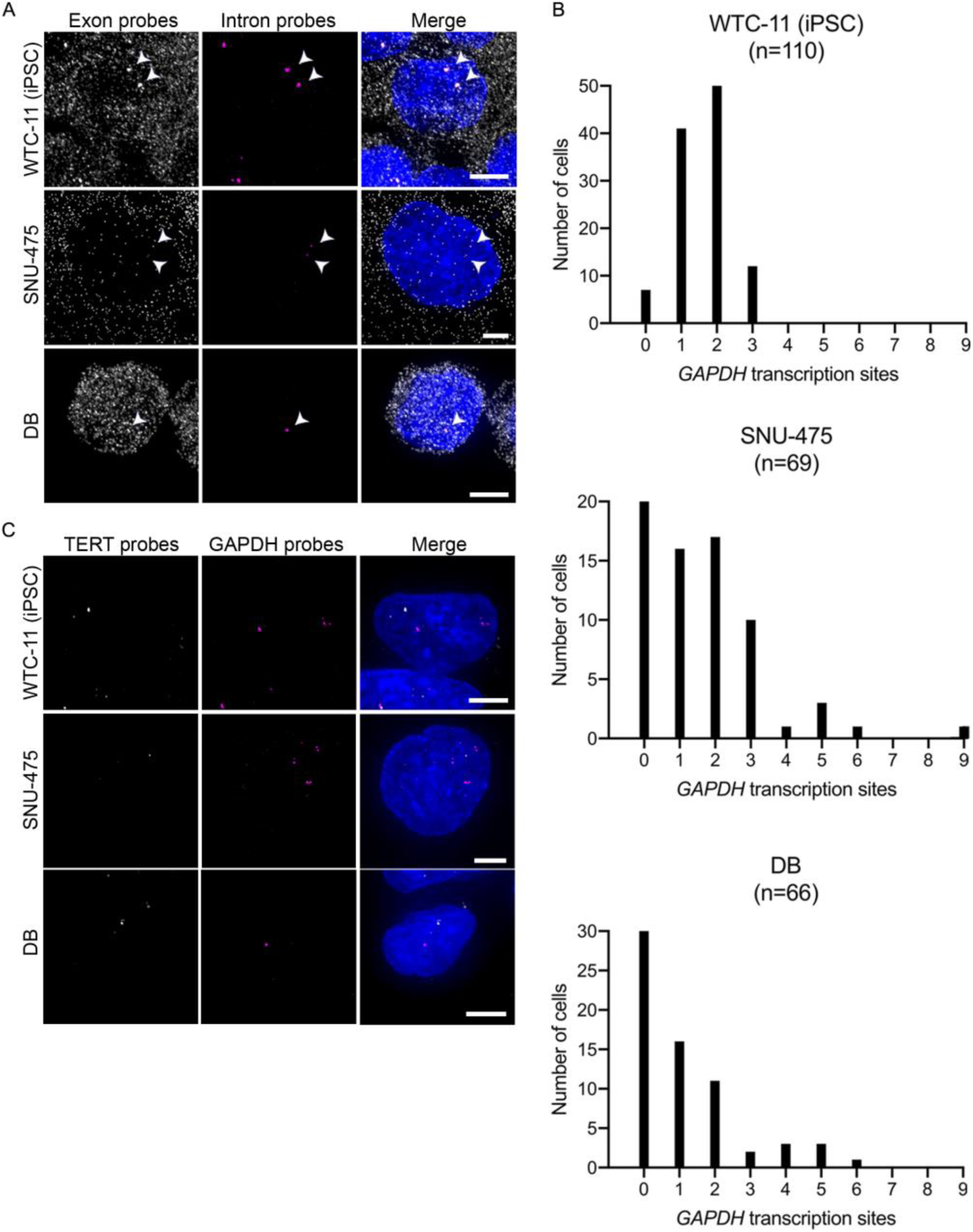
GAPDH smFISH validates the specificity of the smFISH assay. (A) Representative single-cell maximum intensity projections of GAPDH exon and intron smFISH. Arrowheads indicate co-localization of exon and intron signal consistent with active transcription sites in iPSCs (line WTC-11) and two cancer cell lines that express relatively low levels of TERT (SNU-475 and DB). (B) Histograms quantifying cell-to-cell variation in number of *GAPDH* transcription sites (number of co-localized exon-intron spots per nucleus) in indicated cell lines. (C) GAPDH intron and TERT intron probes do not colocalize within the same cells. Data information: GAPDH exon probes are shown in gray, GAPDH intron probes in magenta, TERT intron probes in gray (**c** only), and DAPI in blue. Scale bars are 5 µm.

**Figure EV4.**
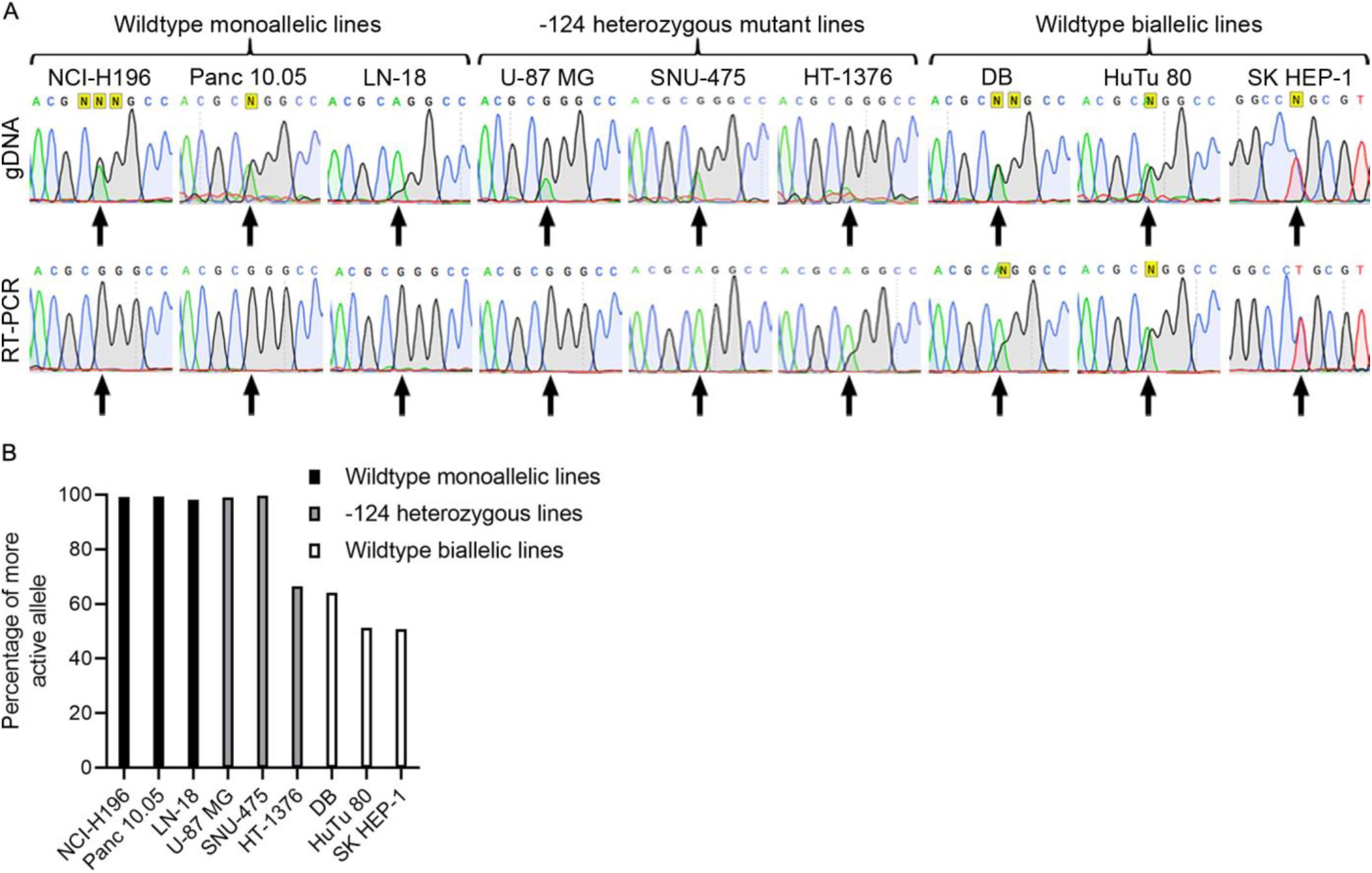
Classification of cancer cells as monoallelic or biallelic *TERT* expressers. (A) gDNA and RT-PCR sequencing of *TERT* exon 2 SNP. (B) Quantification of the more active allele based entirely on RT-PCR results shown in (A) agrees in most cases with the previous classification of these lines as monoallelic (100% one allele) or biallelic (~50% one allele).

**Figure EV5.**
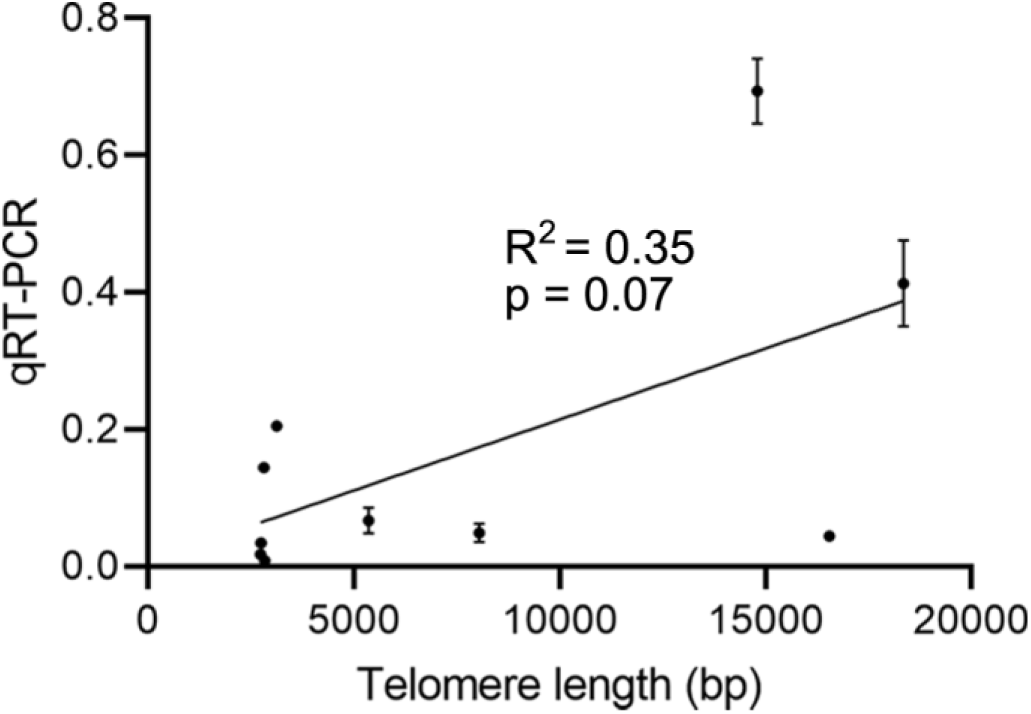
Telomere length, as quantified from the TRF assay shown in Fig. 4, has little correlation with TERT RNA levels, as measured via qRT-PCR, for TERT-expressing cancer lines and line WTC-11 (iPSC). Error bars represent standard error of the mean; n = 3 independent measurements.

